# Massive parallel variant characterization identifies *NUDT15* alleles associated with thiopurine toxicity

**DOI:** 10.1101/740837

**Authors:** Chase C. Suiter, Takaya Moriyama, Kenneth A. Matreyek, Wentao Yang, Emma Rose Scaletti, Rina Nishii, Wenjian Yang, Keito Hoshitsuki, Minu Singh, Amita Trehan, Chris Parish, Colton Smith, Deepa Bhojwani, Liz YP Yuen, Chi-kong Li, Chak-ho Li, Yung-li Yang, Gareth J Walker, James R Goodhand, Nicholas A Kennedy, Federico Antillon Klussmann, Smita Bhatia, Mary V. Relling, Motohiro Kato, Hiroki Hori, Prateek Bhatia, Tariq Ahmad, Allen E. J. Yoeh, Pål Stenmark, Douglas M. Fowler, Jun J. Yang

**Affiliations:** Department of Pharmaceutical Sciences, St. Jude Children’s Research Hospital, Memphis, TN, USA; Department of Genome Sciences, University of Washington, Seattle, WA, USA; Department of Biochemistry and Biophysics, Arrhenius Laboratories for Natural Sciences, Stockholm University, Sweden; Department of Experimental Medical Science, Lund University, Lund, Sweden; Department of Pediatrics, Advanced Pediatrics Centre, PGIMER, Chandigarh, India; Department of Pediatrics, Children’s Hospital of Los Angeles, Los Angeles, California, USA; Department of Pathology, Hong Kong Children’s Hospital, Hong Kong; Department of Paediatrics, The Chinese University of Hong Kong, Hong Kong; Department of Paediatrics and Adolescent Medicine, Tuen Mun Hospital, Hong Kong; Department of Laboratory Medicine and Pediatrics, National Taiwan University Hospital, College of Medicine, National Taiwan University, Taipei, Taiwan; Department of Gastroenterology, Royal Devon and Exeter Hospital NHS Foundation Trust, Exeter, England; IBD Pharmacogenetics Group, University of Exeter, Exeter, England; Unidad Nacional de Oncología Pediátrica, Guatemala City, Guatemala, Francisco Marroquin Medical School, Guatemala City, Guatemala; Division of Pediatric Hematology/Oncology, Institute for Cancer Outcomes and Survivorship, School of Medicine, University of Alabama at Birmingham; Department of Pediatric Hematology and Oncology Research, National Center for Child Health and Development, Tokyo, Japan; Department of Pediatrics, Mie University Graduate School of Medicine, Mie, Japan; Centre for Translational Research in Acute Leukaemia, Department of Paediatrics, Yong Loo Lin School of Medicine, National University of Singapore, Singapore; Cancer Science Institute of Singapore, National University of Singapore, Singapore; Department of Bioengineering, University of Washington, Seattle, WA, USA; Genetic Networks Program, CIFAR, Toronto, ON, Canada

## Abstract

As a prototype of genomics-guided precision medicine, individualized thiopurine dosing based on pharmacogenetics is a highly effective way to mitigate hematopoietic toxicity of this class of drugs. Recently, *NUDT15* deficiency was identified as a novel genetic cause of thiopurine toxicity, and *NUDT15*-informed preemptive dose reduction is quickly adopted in clinical settings. To exhaustively identify pharmacogenetic variants in this gene, we developed massively parallel NUDT15 function assays to determine variants’ effect on protein abundance and thiopurine cytotoxicity. Of the 3,097 possible missense variants, we characterized the abundance of 2,922 variants and found 54 hotspot residues at which variants resulted in complete loss of protein stability. Analyzing 2,935 variants in the thiopurine cytotoxicity-based assay, we identified 17 additional residues where variants altered NUDT15 activity without affecting protein stability. We identified structural elements key to NUDT15 stability and/or catalytical activity with single amino-acid resolution. Functional effects for *NUDT15* variants accurately predicted toxicity risk alleles in 2,398 patients treated with thiopurines, with 100% sensitivity and specificity, in contrast with poor performance of bioinformatic prediction algorithms. In conclusion, our massively parallel variant function assays identified 1,103 deleterious *NUDT15* variants, providing a comprehensive reference of variant function and vastly improving the ability to implement pharmacogenetics-guided thiopurine treatment individualization.

## Introduction

Thiopurines (e.g., mercaptopurine [MP], 6-thioguanine [TG] and azathioprine [AZA]) are important antimetabolite drugs with diverse clinical indications. For example, as a potent anti-leukemia agent, MP-based maintenance therapy is arguably one of the most critical components of the curative treatment regimen for acute lymphoblastic leukemia (ALL) in children and adults^1–5^. Thiopurines are also commonly used as immunosuppressive agents such as rheumatoid arthritis and inflammatory bowel diseases (IBD)^6–8^. Extensive intracellular metabolism of thiopurine prodrugs is required for therapeutic efficacy across diseases. Particularly for their cytotoxic effects, thiopurines need to be converted to thioguanosine triphosphate (TGTP) which is incorporated into DNA to form DNA-TG, triggering futile DNA damage repair and ultimately apoptosis^9–12^.

Genetic variations in genes encoding thiopurine metabolizing enzymes can directly influence drug toxicity and anti-leukemic efficacy^13–16^. For example, genetic polymorphisms in thiopurine methyltransferase *TPMT* have been linked to susceptibility to thiopurine-induced hematopoietic toxicity in patients, and preemptive *TPMT* genotype-guided dosing is one of the first examples of genetics-based precision medicine in cancer^17, 18^. More recently, we and others identified inherited NUDT15 deficiency gene as a major genetic cause for thiopurine intolerance in ALL and IBD patients, most frequently in patients of Asian and Hispanic descent^19–21^. *NUDT15* encodes a nucleotide diphosphatase that inactivates TGTP by converting it to thioguanosine monophosphate (TGMP). Thus, NUDT15 functions as a negative regulator of intracellular TGTP, with loss-of-function *NUDT15* variants leading to accumulation of DNA-TG and increased cytotoxicity^21, 22^.

With the clinical implementation of *NUDT15*-guided thiopurine dosing^18^, this gene is frequently sequenced in thiopurine-treated patients and novel variants are regularly discovered^23^. However, functional consequences of these novel *NUDT15* variants remain largely uncharacterized, thus hampering the implementation of individualized thiopurine therapy. In fact, accurate prediction of the phenotypic effect of genetic variation is a particularly formidable challenge in pharmacogenetics^24^. Bioinformatic prediction algorithms assess the essentiality of a given gene (or genetic variant) and thus their impact on fitness. This assumption is valid for genetic variants related to disease pathogenesis, e.g., deleterious variants in tumor suppressor genes would be under negative selection during evolution^25, 26^. However, genetic variants in pharmacogenes are not subjected to purifying selection because many of them are only involved in xenobiotic metabolism and non-essential in normal physiological conditions^27^. Therefore, experimental characterization is needed to determine the function of pharmacogene variants, but traditional methods are low throughput, laborious, and outpaced by the rate at which novel variants are discovered.

To address this challenge, we utilized massively parallel variant function assays to exhaustively identify *NUDT15* variants that alter protein abundance and/or thiopurine sensitivity. In this systematic screen, we scored 91.8% of the 3,097 possible missense variants in *NUDT15*, of which 1,103 variants resulted in loss of activity. Our high-resolution variant-activity map pinpointed structural features essential for NUDT15 activity. Applying the functional effects-based *NUDT15* variant classification, we accurately predicted thiopurine toxicity risk variants identified in 2,398 patients with 100% sensitivity and specificity.

## Results

To comprehensively characterize *NUDT15* variant function, we first constructed a site-saturated mutagenesis library of 3,077 missense variants in this gene, representing 99.3% of all possible amino acid change across the 163 residues in this protein (Fig. 1). This library of variants was then introduced into an engineered HEK293T landing pad cell line at a defined genomic locus via Bxb1 mediated recombination^28, 29^. Each cell in the library harbored only a single *NUDT15* variant, enabling multiplex evaluation of variant function by measuring cellular phenotypes of interest. Additionally, each variant NUDT15 sequence was tagged with a set of random barcodes (at an average of 14, ranging from 1 to 54). Barcodes were measured individually in function assays, therefore representing independent observations of each variant. To functionally characterize *NUDT15* variants, we developed two massively parallel assays that measure the abundance of variant protein and variants’ effect on thiopurine cytotoxicity, respectively.

**Figure 1.**
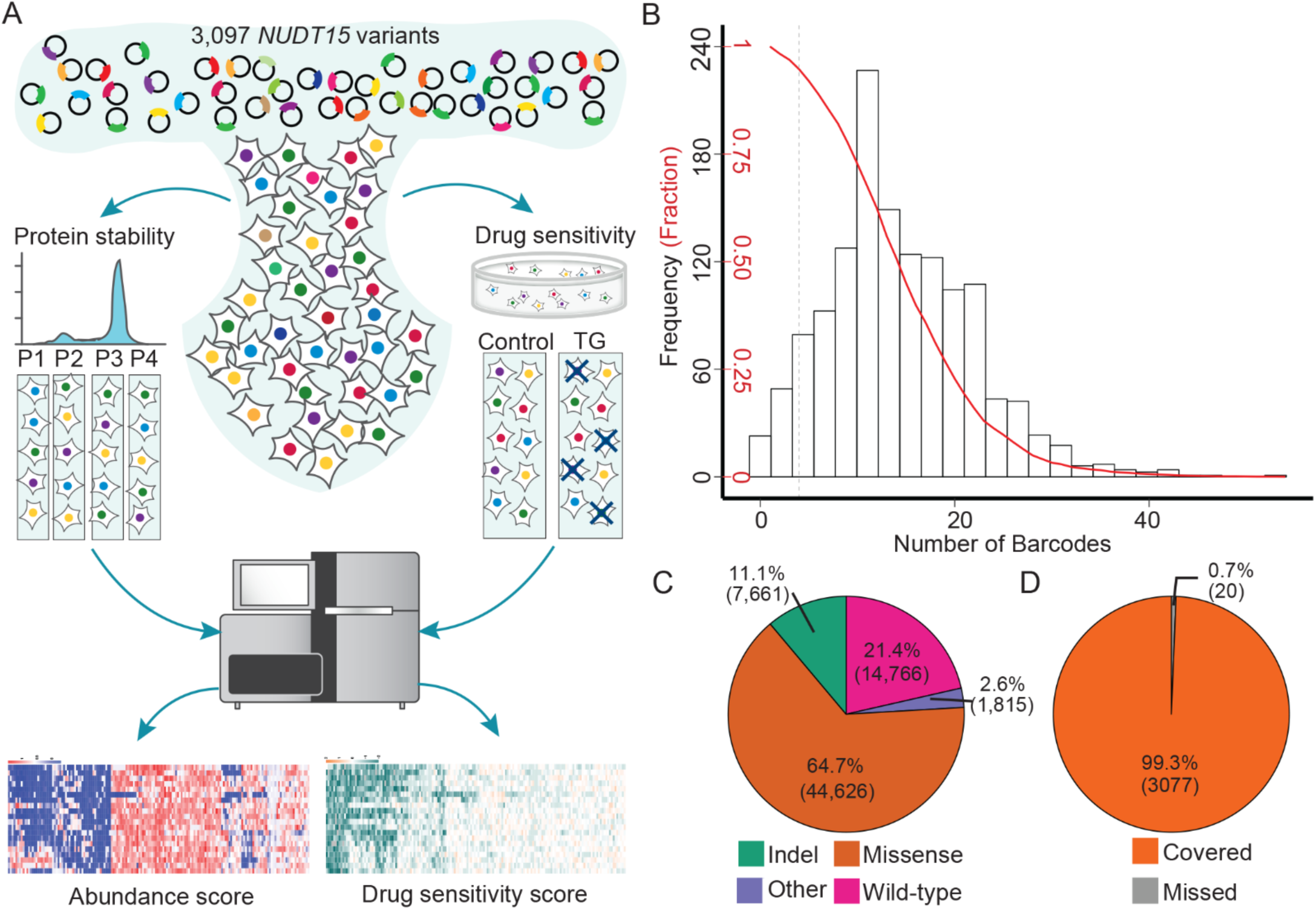
Massive parallel characterization of variant function in *NUDT15*. **A**) *NUDT15* variant library was introduced to HEK293T landing pad cells such that each cell would express only one copy of a specific *NUDT15* variant. Cells were then subjected to two types of phenotyping to determine the effect of a given variant on 1) intracellular NUDT15 protein abundance (measured as the fluorescence intensity of the NUDT15-EGFP fusion protein), or 2) thiopurine cytotoxicity *in vitro* (measured as the frequency of variant-expressing cells after TG treatment). To estimate a variant’s “abundance score”, cells were flow sorted into four groups with decreasing fusion protein fluorescence; and variants over-represented in low fluorescence group were associated with NUDT15 protein stability. To assign variant “drug sensitivity score”, each variant was enumerated in library-transfected cells at baseline and after TG treatment *in vitro*; deleterious variants resulted in low NUDT15 activity and rendered cells sensitive to thiopurine, thus became under-represented after drug exposure. **B**) Distribution of number of barcodes in the *NUDT15* library is shown along with the cumulative fraction of barcoded variants (red line). The library included total 68,868 unique barcodes, each of which was assigned to a specific variant (median 14 barcodes per variant [ranging from 1 to 54]). **C**) In the *NUDT15* variant library, there are 44,626 unique barcodes linked to missense variants, as determined by long-read PacBio sequencing. All together, exhaustive mutagenesis of *NUDT15* gene generated 3,077 missense variants across 163 amino acids, representing 99.3% of all possible missense variations. Abbreviation. TG, 6-thioguanine.

We elected to focus on NUDT15 protein abundance as the functional endpoint for our first massively parallel assay because the majority of clinically actionable *NUDT15* variants known thus far exhibit significant decrease in protein thermostability^21, 30^. In this high throughput screen, individual *NUDT15* variants were fused to the *EGFP* gene, with the fluorescence intensity from the fusion protein as a proxy marker of the steady-state abundance of the variant^29^. As shown in Supplementary Fig. 1A, wildtype (WT) NUDT15 tagged with EGFP at the C-terminus resulted in a fluorescence signal that was easily detectable by flow cytometry. By contrast, expression of the known low-stability variant Arg139Cys fused in the same manner to EGFP resulted in an approximate 3 fold reduction in EFGP signal. Applying this to the *NUDT15* mutagenesis library, we sought to quantify intracellular abundance of all possible variants in parallel. Upon transduction, the population of cells expressing the *NUDT15* library exhibited a left-skewed distribution of normalized EGFP signal, with a predominant peak encompassing WT-expressing cells and a thin tail of cells expressing presumptive destabilizing variants (including Arg139Cys, Supplementary Fig. 1B). Cells were sorted into four equally-populated bins with decreasing fluorescence signal, representing variants with decreasing levels of abundance. Subsequent high-throughput sequencing of cells in each bin allowed the calculation of individual variant frequencies, from which an abundance score was empirically determined for every variant in the library (ranging from −0.31 to 1.47, Fig. 2A, Supplementary Fig. 2A and 2B). After excluding 154 variants due to low quality sequencing, we estimated 2,923 abundance scores representing 94.4% of all possible missense variants in this gene. On average, each variant was measured 14 times (i.e., 14 independent barcodes per variant, Supplementary Fig. 2A). Notably, 735 variants exhibited an abundance score lower than that of the known toxicity risk variant Arg139Cys, suggesting they had severe effects on *NUDT15* abundance and possibly thiopurine metabolism. To validate the results from the high throughput screen, we selected 14 *NUDT15* variants with a wide range of abundance scores for assessment using orthogonal methods. Their abundance scores from the library screen were highly correlated with the EGFP signal of HEK293T cells individually transfected with each variant (R^2^ = 0.96, P < 2.2 x 10^-16^, Supplementary Fig. 3A). We also expressed and purified these 14 NUDT15 variant proteins in *E. coli* and subjected them to a thermal stability assay. Variant T_m_ values normalized to WT protein were strongly correlated with abundance scores (R^2^ = 0.72, P = 6.0 x 10^-5^, Supplementary Fig. 3B). Therefore, the high throughput intracellular abundance screen accurately predicted individual variant protein stability. This massively parallel variant characterization also revealed biochemical determinants of NUDT15 stability. The active NUDT15 enzyme complex consists of two identical monomers, each with a highly conserved NUDIX motif and also a NUDIX fold^22^. The NUDIX motif (GX_5_EX_7_REUXEEXGU) contains the catalytic amino acids, whereas the NUDIX fold is formed by a β sheet with α helices on each side acids. Examining all 19 variations at each residue, we identified 54 hotspot positions at which genetic variants uniformly gave rise to unstable NUDT15 proteins compared to WT (Fig. 2B). These hotspot residues are unevenly distributed: variations at positions with small side chain amino acid (e.g., alanine substitution) were tolerated in general, whereas changes affecting the hydrophobic (valine, proline, leucine, isoleucine and phenylalanine) or nonpolar aliphatic amino acids (methionine) resulted in large decreases in NUDT15 stability (Fig. 2A, Supplementary Fig. 2C). Changes to proline were also poorly tolerated (Fig. 2A), because of the unique backbone geometry imposed by its side chain. Some secondary structures were found to be more vulnerable to substitution relative to flexible loop regions. Of the 54 hotspot residues, 68.5 % were located in secondary structure elements (α-helices, β-sheets and 3_10_ helices, Fig. 2C), especially beta-strands β1, β5 and β6. For example, our screen identified a cluster of hotspots in the β1 strand, with the valine residues within this glycine/valine repeat consistently vulnerable to genetic variation (Fig. 2D). In the crystal structure, these valine side chains strongly interact with the α1 helix to maintain the NUDIX domain conformation. In fact, a number of indel variants in this region also lead to unstable NUDT15 protein and are associated with thiopurine toxicity in patients^21, 31^. Finally, we observed a significant over-representation of hotspot residues at the dimer interface of NUDT15, with 13 (30.9%) of 42 amino acid in this region extremely vulnerable to genetic variation in the abundance-based scree (Fig. 2E). This result strongly argued for the importance of inter-monomer interaction in maintaining the overall stability of NUDT15 protein.

**Figure 2.**
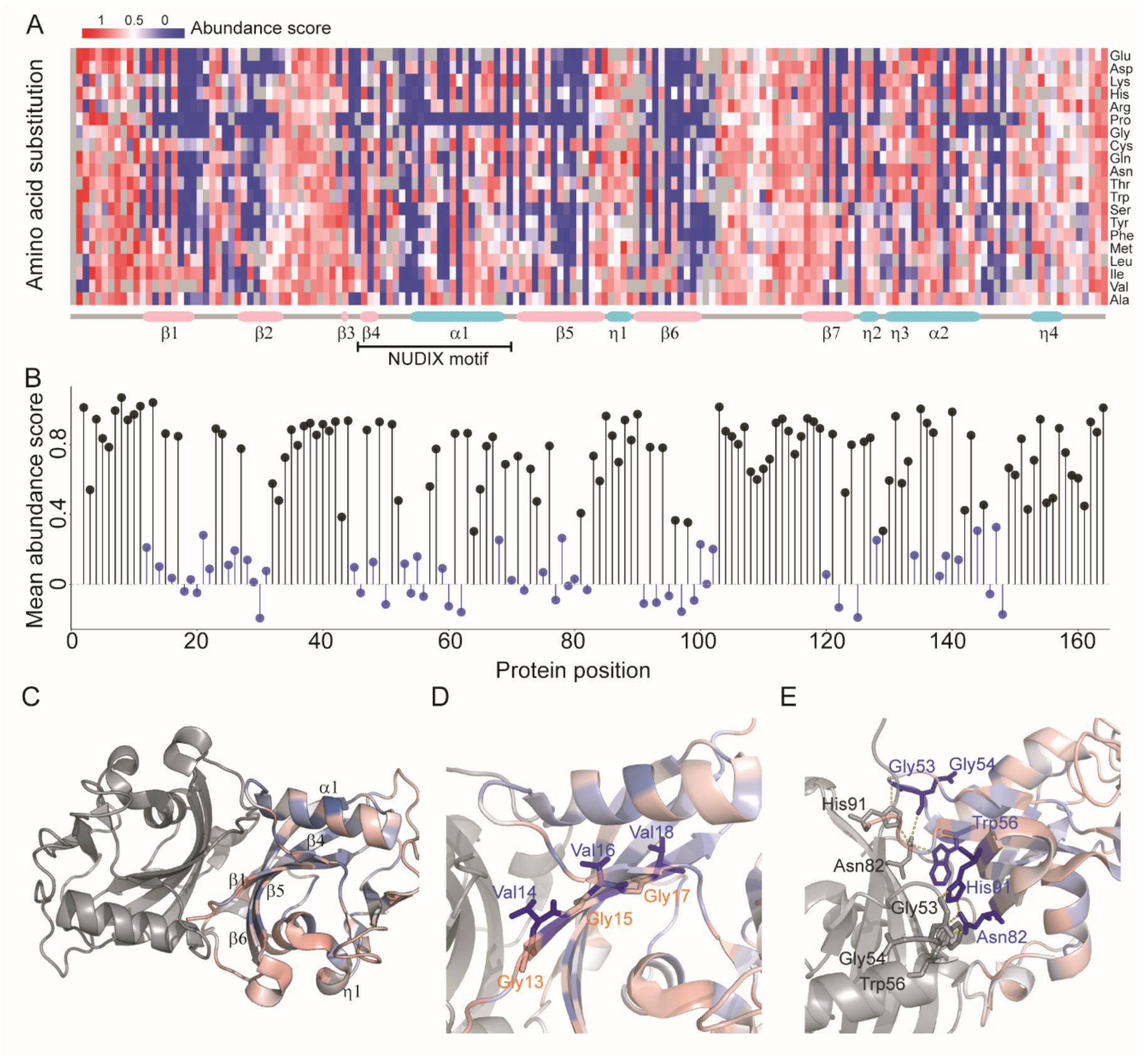
Effects of genetic variants on NUDT15 protein abundance. **A**) NUDT15 abundance score was assigned to 2,922 variants, as plotted in the heatmap. Each column represents an amino acid residue in NUDT15 protein [from 1 to 164], and rows indicate all 19 possible missense changes from the wildtype sequence at this position. Red to blue denotes high and low protein abundance, respectively. Secondary structures are schematically indicated below the heatmap. **B**) An average abundance score was calculated for each NUDT15 residue (mean of 19 variants) and positions at which variants consistently encode low abundance protein were considered as hotspots (54 in total) and highlighted in blue (see **Methods**). **C**) Structural analysis of hotspot residues identified features critical to NUDT15 protein stability. In the 3-dimentional structure (PDB: 5LPG), NUDT15 protein is shown as a homodimer with each subunit in either grey or gradient colors representing mean abundance score respectively. α-helix and β-sheets distal to the NUDIX motif (valines at 14, 16 and 18 in GlyVal repeats in Panel **D**) and residues at the interface between two monomers (Gly53, Gly54, Trp56, Asn82 and His91 in Panel **E**) are particularly enriched with hotspot residues.

Although abundance-based screening identified a significant number of deleterious *NUDT15* variants, we postulate that there are other mechanisms by which genetic variants cause loss of function without affecting protein abundance (e.g., alterations of substrate binding). Thus, we performed a secondary massively parallel screen to directly examine the effects of *NUDT15* variants on thiopurine cytotoxicity *in vitro*. In this assay, HEK293T cells harboring the *NUDT15* variant library were treated with 3 μM TG for 6 days and high throughput sequencing was again employed to identify the frequency of each variant prior to drug treatment as well as in cells surviving thiopurine exposure. Because cells expressing loss-of-function NUDT15 variants were more susceptible to thiopurine-induced apoptosis, we estimated a drug sensitivity score for each variant based on its decrease in frequency after drug treatment (Fig. 1A). We successfully evaluated 2,935 variants (94.7 % of the library) in this cytotoxicity assay, with an average of 14 independent observations per variant (Fig. 3A, Supplementary Fig. 4A, and 4B).

**Figure 3.**
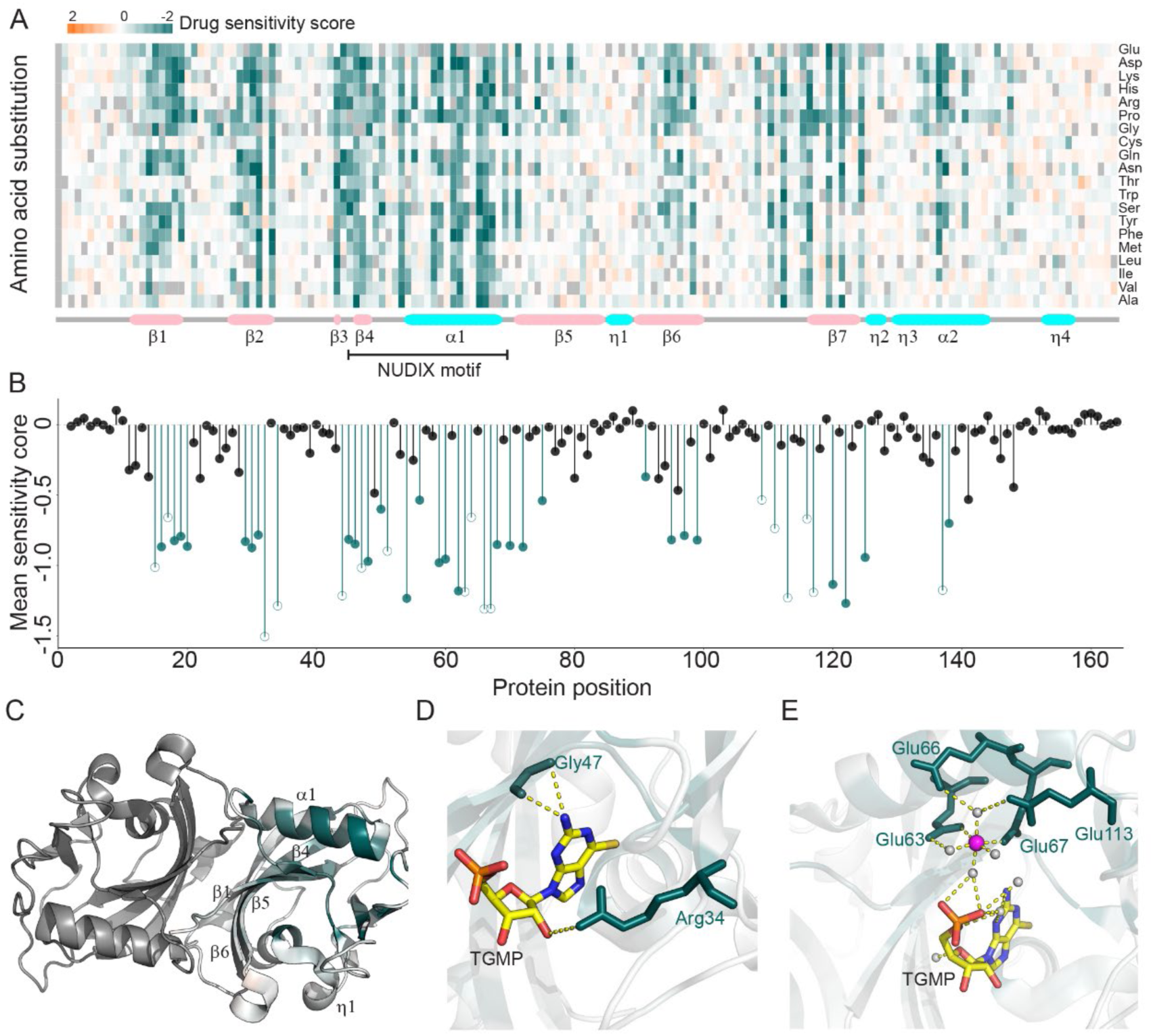
Effects of *NUDT15* variants on thiopurine cytotoxicity. **A**) Drug sensitivity score was assigned for 2,934 variants, as plotted in the heatmap. Each column represents an amino acid residue in NUDT15 protein [from 1 to 164], and rows indicate all 19 possible missense changes from the wildtype sequence at this position. Red and green denotes wildtype-like and damaging *NUDT15* variants, respectively. Secondary structures are schematically indicated below the heatmap. **B**) An average drug sensitivity score was calculated for each NUDT15 residue (mean of 19 variants) and positions at which variants consistently result in increased thiopurine sensitivity were considered as hotspots and highlighted in green (see **Methods**). Among the 45 hotspot residues, 28 were also identified in abundance-based screen (filled circle), whereas 17 amino acid positions were uniquely sensitive to variation in the drug sensitivity-based assay (open circle). **C**) Structural analysis of hotspot residues identified variants critical for thiopurine cytotoxicity. In the 3-dimentional structure (PDB: 5LPG), NUDT15 protein is shown as a homodimer with each subunit in either grey or gradient colors representing mean drug sensitivity score, respectively. In particular, residues involved in direct substrate interaction (Arg34 and Gly47 in Panel **D**) and those interacting with the magnesium ion (magenta) or water (gray) (Glu63, Gllu66, Glu67 and Glu113 in Panel **E**) strongly influence catalytic activity without affecting protein stability. Abbreviations. TGMP, thioguanosine monophosphate.

Similarly, 45 residues were classified as hotspots because all 19 possible variations at each position consistently led to increased thiopurine sensitivity compared to WT (Fig. 3B and Supplementary Fig. 4C). In line with the results from the abundance-based screen, a majority of loss-of-function variants identified from the drug sensitivity-based assay were also located in secondary structure elements. Ten hotspot residues (Gly32, Arg34, Gly47, Gly48, Glu54, Arg62, Glu63, Glu66, Glu67 and Glu113) were either in close proximity to or located within the highly conserved NUDIX motif (α1 helix, Fig. 3C)^22^. This motif contains residues responsible for the coordination of magnesium and water molecules and is directly involved in TGTP hydrolysis^30^. Interestingly, of the ten NUDIX domain residues, four (Gly32, Gly48, Glu54 and Arg62) were also associated with low abundance. In contrast, *NUDT15* variants affecting Arg34 and Gly47 dramatically altered thiopurine sensitivity with minimal effects on protein stability (Fig. 3D). This is also true for residues involved in magnesium coordination (Glu63 and Glu67) or interaction with magnesium-coordinating water molecules (Glu66 and Glu113, Fig. 3E).

To summarize the effects on abundance and drug sensitivity, we scaled both scores from zero to one, and for each variant selected the lower of the two as the final NUDT15 activity score (Fig. 4A, Supplementary Fig. 5A). We defined variants below 0.4 as damaging after modeling the bimodal distribution of activity score, representing 1,103 (35.6%) of all 2,844 variants. Our systematic experimental characterization also allowed for a direct comparison with damaging effects predicted *in silico* by a number of different algorithms: the combined annotation dependent depletion (CADD) score, the rare exome variant ensemble learner (REVEL) score (Fig. 4B and 4C), polymorphism phenotyping (PolyPhen2), and sorting intolerant from tolerant (SIFT) (Supplementary Fig. 5B and 5C). With CADD score > 20 as the criterion for damaging variants^25, 32^, 561 variants were predicted as loss-of-activity, of which 308 exhibited a high NUDT15 activity score (i.e., > 0.4), with a false positive rate of 54.9%. Conversely, of 276 CADD-predicted benign variants, 12 had low NUDT15 activity score, with a false negative rate of 4.3%. Similarly, when we performed this analysis using the REVEL prediction (>0.5 as damaging^26^), we observed a false positivity and false negativity rate of 28.3% and 25.9%, respectively.

**Figure 4.**
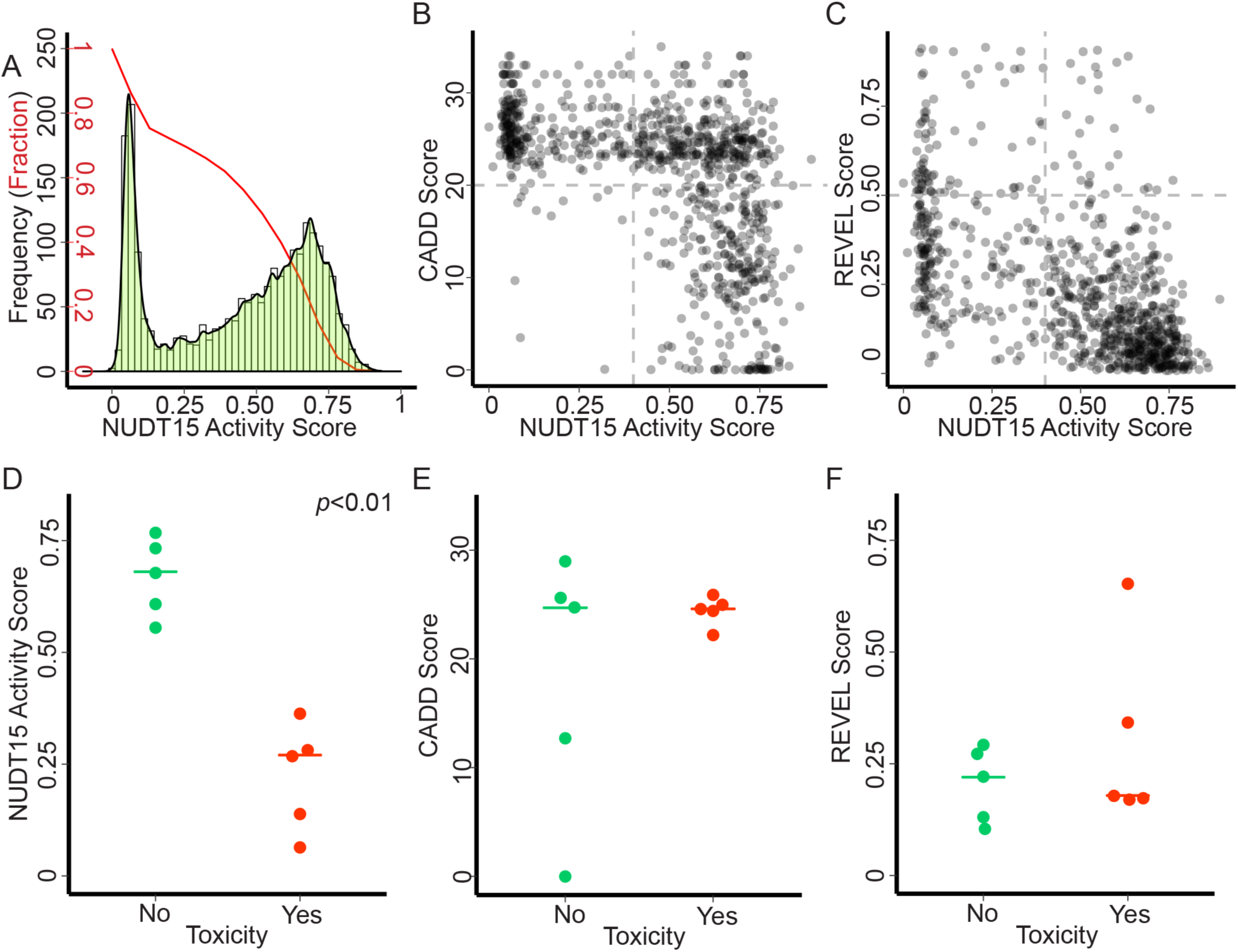
NUDT15 activity score predicted clinical thiopurine toxicity. **A**) Combining abundance score and drug sensitivity score, we assigned a final NUDT15 activity score for 2,844 variants, whose distribution is plotted along with the cumulative fraction of variants (red line). Comparison of experimentally-determined NUDT15 activity score with effects predicted by bioinformatic algorithms (i.e., panels **B and C** for CADD and REVEL scores, respectively). CADD score (>20 as damaging) and REVEL score (> 0.5 as damaging) were available for 837 missense variants. Horizontal and vertical dashed lines represent cutoff for NUDT15 activity score and CADD/REVEL scores, respectively. *NUDT15* variants were identified by sequencing in 2,398 patients exposed to thiopurine therapy and were classified as toxicity-related or non-toxicity related (see **Methods**). NUDT15 activity (**D**), CADD (**E**), or REVEL scores (**F**) were plotted for risk vs benign variants, with P value estimated by using the Mann-Whitney-Wilcoxon test.

To apply our functional effects-based variant classification, we sought to identify *NUDT15* variants in patients treated with thiopurine drugs and ask if activity score could predict pharmacogenetic variants associated with toxicity. In 2,398 subjects, we identified a total of 10 missense coding variants, of which six were novel and exceedingly rare. Cases carrying *TPMT* variants were excluded for further analyses. Five variants were associated with hematopoietic toxicity secondary to thiopurines (Lys33Glu, Arg34Thr, Val75Gly, Arg139Cys and Arg139His), whereas five were not (Gln6Glu, Arg11Gln, Val18Ile, Ser83Tyr and Val93lle, Supplementary Table 1). Activity score averaged at 0.23 for five toxicity variants (range from 0.061 to 0.38), significantly lower than that of variants not linked to toxicity (mean of 0.69, ranging from 0.55 to 0.81, P = 0.0079, Fig. 4D). With 0.4 as the activity score cutoff, we estimated the sensitivity and specificity both at 100%, based on the prediction of these 10 variants. In contrast, CADD (>20) or REVEL (>0.5) scores did not predict variants’ effect on thiopurine toxicity, with sensitivity and specificity at 100%, 40%, 20%, and 100%, respectively (Fig. 4E and 4F). The receiver operating characteristic curve analysis projected a probability of 1.0 of accurate prediction of risk variants using activity score (area under the curve statistic, AUC), whereas AUC was 0.48 and 0.64 for the CADD and REVEL-based prediction (Supplementary Fig. 6).

Querying the publicly available gnomAD database of whole genome/exome sequencing of 141,456 individuals, we identified 108 *NUDT15* missense alleles observed in human with the population frequency ranging from 4 in 100,000 to 2:100 (Fig. 5A). Damaging variants were detected in all populations regardless of ancestry, and the average NUDT15 variant activity score did not differ by race or ethnicity (Fig. 5B). Of 8,871 individuals in gnomAD with *NUDT15* variation, 8,125 (91.6%) have a damaging variant and thus are at risk of thiopurine toxicity (7,399 were attributable to the Arg139Cys variant, Fig. 5C).

**Figure 5.**
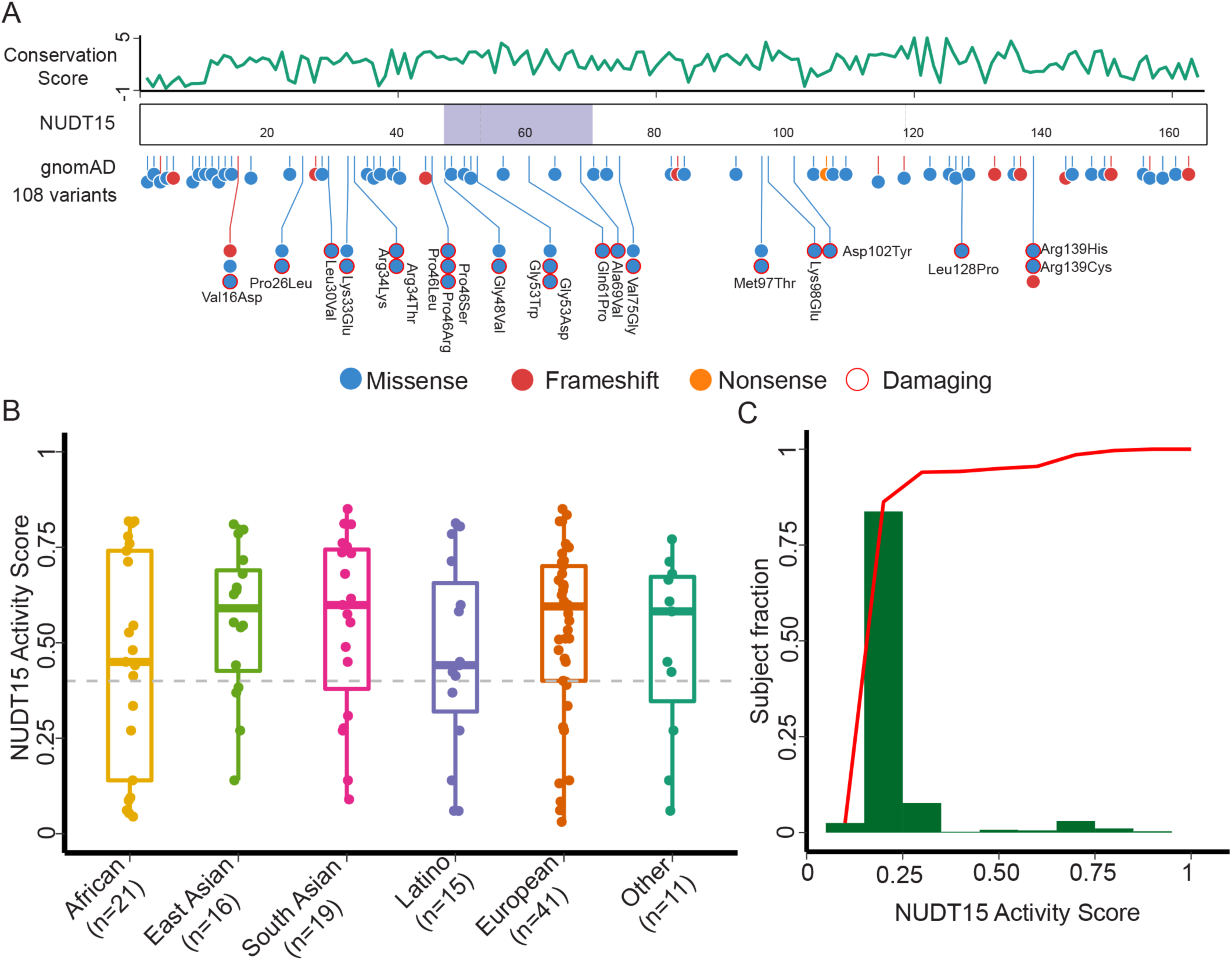
Population distribution of *NUDT15* variants in humans. **A**) 108 *NUDT15* variants are identified in whole genome or whole exome seq data of 141,456 individuals in the gnomAD database (https://gnomad.broadinstitute.org/). In the lollipop plot, each circle represents a *NUDT1*5 variant (blue, red, or orange for missense, frameshift, or nonsense, respectively). Damaging *NUDT15* variants are annotated with the exact amino acid change and open red circles. Average PhastCons scores of trinucleotide are shown in green line at the top for each amino acid residue to indicate cross-species conservation. The Nudix domain is highlighted in purple. **B**) Activity score is plotted for NUDT15 variants observed in each of 5 major race/ethnic groups in the gnomAD cohort. Damaging variants (NUDT15 activity score < 0.40, dashed line) were present across populations. Numbers of variants identified in each population are shown in parenthesis. **C**) 91.6% of individuals affected by *NUDT15* polymorphism carry a variant that results in significant loss of NUDT15 activity. This is in part explained by the common variant Arg139Cys (with an activity score of 0.14).

## Discussion

Accurate annotation of the phenotypic effect conferred by genetic variation is of critical importance for the implementation of genomics-guided precision medicine^24^. This has become a pressing challenge in recent years with the explosive growth of genome sequencing and the sheer number of novel variants that need to be functionally characterized. In fact, 48.8% of variants cataloged in the ClinVar database are considered of “unknown significance”, primarily due to the lack of experimental validation of their functional effects^33^. Moreover, the vast majority of human genetic variants are rare^34^, for which statistical association with clinical phenotypes would be exceedingly difficult to establish. Thus, functional experiments are essential for predicting the clinical consequences of rare genetic variants. To address this, a number of groups have utilized high-throughput genomics platforms to evaluate variant function at scale^24, 29, 35, 36^. Similarly, we report a preemptive large-scale screen of functional variants in *NUDT15* and identified 1,103 loss-of-function variants, and our clinical validation study confirmed the predicted association with thiopurine toxicity. These results provide a comprehensive catalog of all possible missense pharmacogenetic variants in this important drug metabolizing gene, vastly improving the ability to implement genotype-guided treatment individualization.

Scalable functional assays are of particular importance for pharmacogenetic variants because computational predictions lack sufficient accuracy to be relied upon clinically^37^. For example, comparing *NUDT15* variants experimentally identified as damaging vs those predicted using CADD, we observed that the bioinformatic algorithm classified a substantial proportion of variants as loss-of-function even though they showed no effects on protein stability or thiopurine cytotoxicity (Fig. 4B and 4C). One potential explanation is that most computational prediction methods rely on (to varying degrees) evolutional conservation of genetic variation with presumable effects on human fitness (e.g., tumor suppressors). However, this assumption that deleterious variants would be selected out during evolution is often times invalid for genes involved in drug metabolism because they are not essential for survival. Until the introduction of pharmaceutical agents a few hundred years ago, there was no selection pressure against pharmacogenetic variants and therefore this type of prediction model performs poorly in this context. That said, it is formally plausible that CADD or REVEL identify modes of damaging effects that are not reflected in our two functional assays, or these variants have subtle effects that our assays were not sufficiently sensitive to detect.

There are a number of caveats with our choice of experimental endpoints for *NUDT15* variant characterization. In particular, the thiopurine cytotoxicity-based screen, while successful at determining variants that most severely alter cell sensitivity to this drug (presumably via effects on thiopurine metabolism), is likely limited in its sensitivity for two reasons. One, our screen employed HEK293T cells which are known to lack certain components of the mismatch repair system^38^, and thus are somewhat more resistant to cell cycle arrest than mismatch repair-proficient cell lines. As a result, the high background of thiopurine resistance in HEK293T cells may have masked effects of variants that modestly influence drug sensitivity. Second, we believe that the degree of variant drop-out after thiopurine exposure is highly dependent on the drug concentration used and length of incubation. While we did discover variants causing the most deleterious alterations with probable structural basis, variants with intermediate metabolic activity may be able to metabolize enough drug to persist over the course of our selection.

In conclusion, we report the results of a deep mutational scan of *NUDT15* for identifying pharmacogenetic variants, creating a comprehensive reference of risk alleles to enable preemptive tailored thiopurine therapy. Our findings also point to the critical importance and exciting potential of high throughput variant annotation in pharmacogenes in general.

## Materials and Methods

### *NUDT15* mutagenesis library and the landing pad cellular model

*NUDT15* variant library was synthesized (Twist Biosciences, CA) to exhaustively introduce missense variants across coding region of this gene, followed by the addition of random sequence barcodes, using procedures previously described^29^. This barcoded *NUDT15* library was integrated into the *AAVS1* locus in the HEK293T landing pad cell line such that each cell would express a single *NUDT15* variant fused with EGFP^28, 29^. Cells with successful recombination were identified by flow cytometry (BFP-negative and mCherry-positive), with which *NUDT15* variant characterization was performed subsequently. Primers used for library construction and cloning are listed in Supplementary Table 3. Detailed descriptions of these experiments are provided in **Supplementary Notes**.

### Protein abundance-based screen of *NUDT15* variants

Abundance score of each variant was determined using the VAMP-seq method^29^ (Fig. 1A and **Supplementary Notes**). Briefly, library-expressing HEK293T cells were first sorted into four different bins depending on the level of EGFP normalized to mCherry by flow cytometry. Massive parallel sequencing was then performed to quantify every variant in each of the four bins, from which a variant abundance score was modeled to indicate its intracellular protein abundance (Supplementary Table 2).

### Thiopurine cytotoxicity-based screen of *NUDT15* variants

For thiopurine cytotoxicity-based screen, *NUDT15* library-expressing cells were treated with 3 µM TG or culture media *in vitro* for six days. Cells were then harvested for genomic DNA extraction, and massive parallel sequencing was performed to quantify variant frequency as described above. The final variant-barcode counts table was used as the input for the ABSSeq pipeline^39^ to normalize and identify variants with differential frequency between drug-treated vs non-treated cells (using aFold module). Log_2_ fold change in variant frequency was used to estimate “drug sensitivity score” for each variant (Supplementary Table 2). Each drug treatment had four replicates.

### Hotspot residues and structural analysis

Hotspot analysis was performed to identify residues at which genetic variation caused consistent damaging effects, as follows: for each amino acid residue, we compared the distribution of abundance score or drug sensitivity score of all 19 variant with that of all *NUDT15* variants in the library; those with lower than population average were considered as hotspot residues (P<0.05 using Mann-Whitney-Wilcoxon Test, adjusted by Benjamini & Hochberg approach^40^). t-SNE analysis was also performed to visualize similarity across NUDT15 residues in the effects of their genetic variants, using the Rtsne algorithm^41^.

For the structural analysis, color gradient representing mean abundance score or mean drug sensitivity score was mapped to NUDT15 protein structure (5LPG, http://www.rcsb.org/pdb/home/home.do). A total of 42 interface residues were identified by querying Protein Interfaces, Surfaces and Assemblies (PISA) of the EBI webserver (https://www.ebi.ac.uk/pdbe/pisa/)^42^. Based on these, structural analyses of hotspot residues were performed using PyMOL (version 2.0, Schrödinger, LLC).

### NUDT15 activity score

Abundance score and drug sensitivity score for each variant was first re-scaled to the same range. The smaller value of the two was then assigned as the final “NUDT15 activity score” for each variant (Supplementary Table 2). The cutoff of activity score (0.4) was selected by Ckmeans.1d.dp on the basis of bimodal distribution of the scores^43^.

### Association of *NUDT15* variants with thiopurine toxicity in patients

To identify novel *NUDT15* variants and evaluate their effects on thiopurine toxicity in patients, we ensembled a cohort of 2,398 patients treated with thiopurine for ALL or IBD, including 1,404 subjects in previously published datasets^20, 21, 31, 44, 45^. The ALL cohort included US Children’s Oncology Group trial AALL03N1 (N = 646), Singapore/Malaysia MaSpore ALL 2003 trial (N = 140), Japanese Children’s Cancer Group ALL B-12 (N = 116), Guatemalan LLAG-0707 study (N = 181), Taiwan TPOG ALL study (N = 1), Hong Kong CCCG-ALL2015 trial (N = 132), and the Indian Childhood Collaborative Leukemia Group study (N = 105), for whom targeted sequencing was performed for all 3 *NUDT15* exons. IBD subjects were from the Exeter pharmacogenetic PRED4 study in United Kingdom (N = 1,077), with *NUDT15* variants identified by whole exome sequencing^31^. *TPMT* risk variants (e.g., rs1800462) were also genotyped as previously reported^20, 21, 31, 44, 45^, and cases carrying *TPMT* variants were excluded from further analysis. This study was approved by the respective institutional review boards, and informed consent was obtained from the parents, guardians, and/or patients, as appropriate. Collectively, we identified a total of 10 missense variants (Arg139Cys, Arg139His, Val18Ile, Gln6Glu, Arg11Gln, Lys33Glu, Arg34Thr, Val75Glu, Ser83Tyr, and Val93Ile). To compare the allelic effect across variants, we excluded cases with homozygous or compound-heterozygous *NUDT15* genotype, and the association with toxicity was evaluated based on the difference between heterozygous cases and individuals with WT *NUDT15* (Supplementary Table 1). Details of this analysis are provided in the **Supplementary Notes**.

## Acknowledgements

We thank the patients and parents who participated in the clinical studies included in this report. This work was supported by the US National Institutes of Health (CA021765, GM115279, GM118578, R01CA096670, U10CA098543, U10CA098413, U10CA095861, U10CA180886, U10CA180899), the American Lebanese Syrian Associated Charities of St. Jude Children’s Research Hospital, the V Foundation for Cancer Research, and the Alex’s Lemonade Stand Foundation (T. Moriyama). This work was also supported by the Swedish Research Council and the Swedish Cancer Society (P. Stenmark).

## Supplementary Notes

### Intracellular abundance of the Arg139Cys variant in the landing pad HEK293T cells

Wildtype (WT) *NUDT15-EGFP* was cloned into attB-IRES-mCherry backbone by using NEBuilder HiFi DNA Assembly Cloning Kit (New England BioLabs) according to the manufacture’s instruction. *NUDT15* variant c.C415T (encodes Arg139Cys) was generated via site-directed mutagenesis by using QuikChange Lightning Site-Directed Mutagenesis Kit (Agilent Technologies). The vectors were transfected in a landing pad HEK293T cells for recombination at the *AAVS1* locus, and *NUDT15* expression was induced using 2 μg/mL doxycycline^1, 2^. Recombinant cells expressing WT or Arg139Cys NUDT15 were identified by flow cytometry as BFP-negative and mCherry-positive. NUDT15 protein abundance in these cells was quantified as EGFP/mCherry ratio (Supplementary Fig. 1A).

### *NUDT15* mutagenesis library and the landing pad cellular model

Library construction, cloning, and transfection were performed according to our previously published procedures, with slight modifications^1, 2^. First, a *NUDT15* site-saturation mutagenesis oligo library with flanking attB sequences was purchased from Twist Biosciences (Menlo Park, CA). The oligo library was introduced into SacI and SphI site on the attP-IRES-mCherry backbone by NEBuilder (New England Biolab), with EGFP coding sequenced fused to the C-terminus of NUDT15. An 18-bp barcode was inserted between the stop codon of EGFP and the beginning of the IRES sequence, using T4 ligase. The relationship between of variants and barcodes was established by single molecule real time (SMRT) sequencing. In all subsequent experiments, these barcodes regions would be sequenced to identify individual *NUDT15* variants.

The barcoded *NUDT15* library was integrated into the *AAVS1* locus in the landing pad HEK293T cell line. Cells were transfected with a Bxb1 expression plasmid and the *NUDT15* library on successive days. Cells were cultured for 48 hours to allow for recombination, then *NUDT15* expression was induced using 2 μg/mL doxycycline. Successfully recombined cells were identified by flow cytometry as BFP-negative and mCherry-positive.

### Protein abundance-based screen of *NUDT15* variants

Recombinant cells were binned into four equally populated groups based on their EGFP/mCherry ratio, and sorted library genomic DNA was separately isolated. PCR was performed to amplify barcode regions and to add a unique molecular index. Paired-end 100bp sequencing was performed on Illumina HiSeq 2000. Sequence analyses (reads mapping and alignment, variant calling) were performed in the accordance with the previous study^1^. Sequenced barcodes counts were then assigned to each *NUDT15* variant according to results from SMRT sequencing generated as above. An abundance score for each variant, representing intracellular variant protein abundance, was calculated on the basis of its read counts in each bin, as described previously^1^.

### Bioinformatic prediction of *NUDT15* variant function

Potential damaging effects of *NUDT15* variants were predicted using a variety of bioinformatic algorithms: 837, 837, 837 and 834 variants were included for CADD (v1.4)^3^, REVEL^4^, PolyPhen2^5^ and SIFT^6^ prediction, respectively (Supplementary Table 2). The association of NUDT15 activity score with each bioinformatic prediction was analyzed by using the Spearman (CADD and REVEL) or the Rank Sum test (PolyPhen2 and SIFT), respectively.

### Orthogonal validation of *NUDT15* variant function

14 *NUDT15* variants were selected to validate the accuracy of the high throughput screen using orthogonal assays. First, individual *NUDT15* variants were introduced to the attB-NUDT15-GFP-IRES-mCherry by site directed mutagenesis using an Agilent QuikChange kit. Each variant was individually introduced into HEK293T cells by transfection, and the EGFP/mCherry ratio was quantified by flow cytometry to indicate intracellular NUDT15 protein levels.

In a second assay, variant *NUDT15* was cloned into the pColdII expression vector with an N-terminal His tag via site directed mutagenesis. Variant NUDT15 protein was expressed in *E. Coli*, purified to homogeneity, and then subjected to thermal stability assay using Sypro®Orange (Molecular Probes) to determine T_m_ value (melting temperature), as described previously^7^.

The correlation of individually assessed EGFP/mCherry ratio or T_m_ values with abundance score of these 14 variants were statistically evaluated by using the Spearman test. The GFP/mCherry ratio and T_m_ values were calculated from three independent measurements.

### Association of *NUDT15* variants with thiopurine toxicity in patients

Considering the differences in how thiopurine therapy is given in different diseases, drug toxicity was evaluated based on tolerated dosage (in patients with acute lymphoblastic leukemia [ALL]) or the degree of myelosuppression at a given dosage (in patients with inflammatory bowel diseases [IBD]), as we described previously^7–12^. For patients with ALL, MP dosage was clinically adjusted during the maintenance phase according to a pre-defined range of white blood cell (WBC) count and/or neutrophil count (ANC), and tolerated MP dosage was defined as the average of the daily dosage over at least 14 days after appropriate dose titration. Standard MP dosage and target WBC/ANC account were comparable with slight variation across different ALL treatment protocols included in this study (Supplementary Table 4). For patients with IBD, thiopurine hematopoietic toxicity was evaluated using a multi-step algorithm as described previously^8^, with myelosuppression defined as WBC ≤ 2.5 x 10^9^ cells/l during thiopurine therapy without dose adjustment.

*NUDT15* variants were divided into “rare” or “common”: variants with an allele frequency > 1% in at least one population in the gnomAD cohort (https://gnomad.broadinstitute.org/) are considered as common (Arg139Cys, Arg139His, and Val18Ile), whilst others variants are classified as rare (Gln6Glu, Arg11Gln, Lys33Glu, Arg34Thr, Val75Glu, Ser83Tyr, and Val93Ile). For common variants, we statistically tested the association of genotype with tolerated MP dosage during ALL therapy. Thus, tolerated MP dosage from patients carrying the *NUDT15* variant was compared to those with WT genotype (matched on race or ethnicity), with P value estimated using the linear regression model after adjusting for cohort.

For rare variants, five were identified in patients with ALL (Gln6Glu, Arg11Gln, Ser83Tyr, Val93Ile, and Arg34Thr) and two from the IBD cohort (Lys33Glu and Val75Glu). ALL patients carrying Gln6Glu, Arg11Gln, Ser83Tyr or Val93Ile tolerated MP dosage of 58.3, 50.0, 52.9 or 50.5 mg/m^2^, all of which were within 10% of the average dosage of ancestry-match WT controls from each protocol^7, 9, 11, 12^. By contrast, the MP dosage of two subjects with the rare variant Arg34Thr was 17.9 or 16.4 mg/m^2^, representing 40.3 % and 36.4 % of WT controls, thus classified as “toxicity-related”. Two rare variants identified in the IBD patients (Lys33Glu and Val75Glu) were both classified as “toxicity-related” on the basis of myelosuppression during thiopurine therapy as described above^8^.

The full result of the association analysis is provided in Supplementary Table 1.

---

**Supplementary Fig 1.**
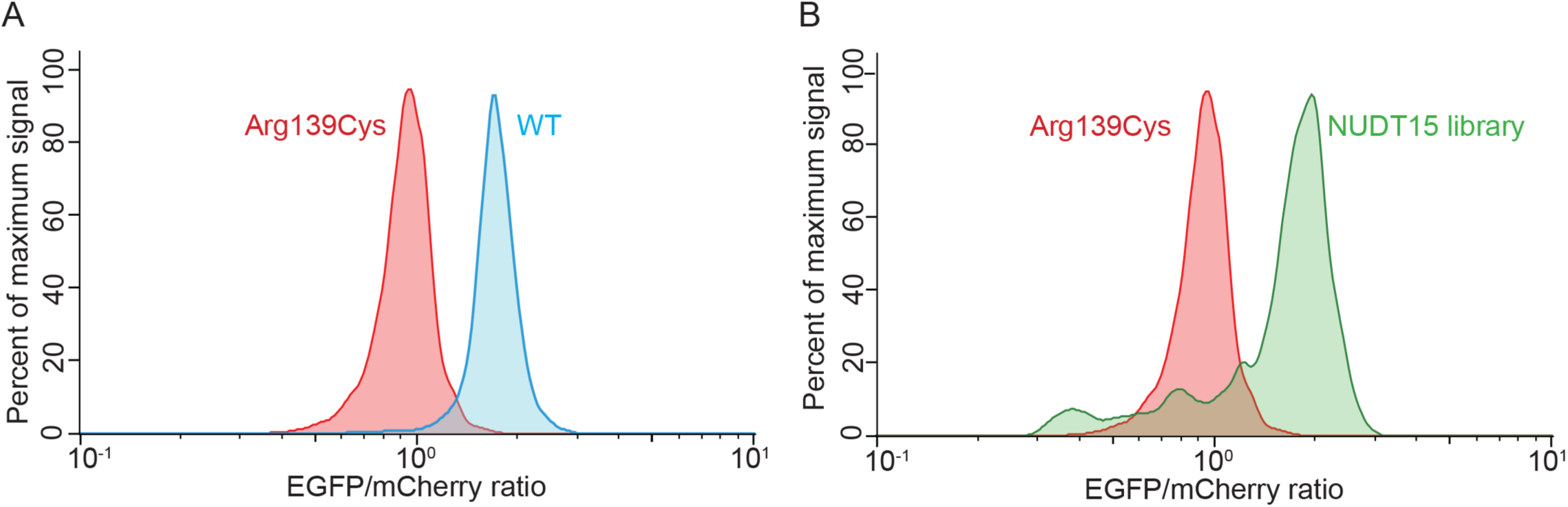
Flow cytometry analysis of HEK293T cells expressing different NUDT15 variants. **A**) Cells were transfected individually with either wildtype (WT) NUDT15 or the Arg139Cys variant (fused with EGFP), and fluorescence was measured by flow cytometry and normalized to mCherry to account for differences in transfection efficiency. **B**) Similarly, fluorescence was measured for cells transfected with the Arg139Cys variant or the entire *NUDT15* variant library.

**Supplementary Fig 2.**
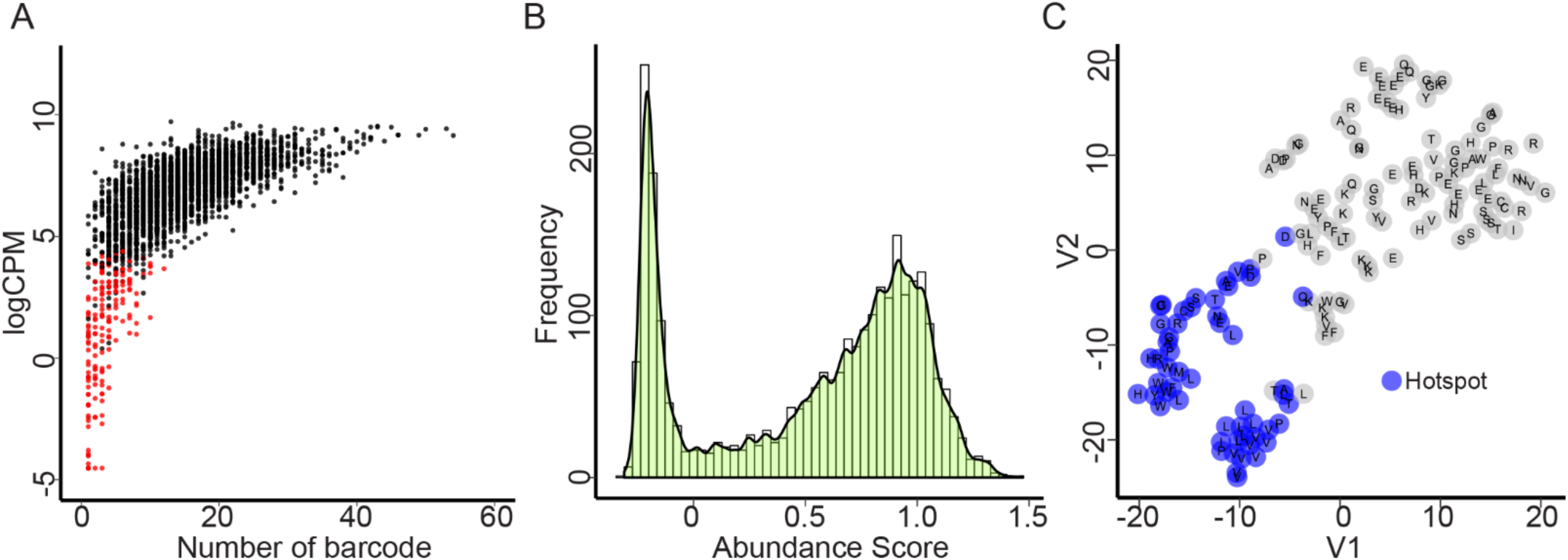
Protein abundance-based functional screen of *NUDT15* variants. **A**) Correlation between sequencing depth and the number of barcodes across 3,097 *NUDT15* variants, with each dot representing a unique variant. After quality control, 154 variants (red dots) were removed and 2,922 variants (black dots) remained for further analysis. **B**) Distribution of variant abundance score. **C**) t-SNE analysis identified residues at which genetic variation has similar effects on NUDT15 protein stability. Each dot represents a unique residue and clustering was on the basis of similarity and differences in the abundance score of 19 variants at each residue. Blue highlight indicates hotspot residues (see **Methods**). Abbreviation. CPM, count per million

**Supplementary Fig 3.**
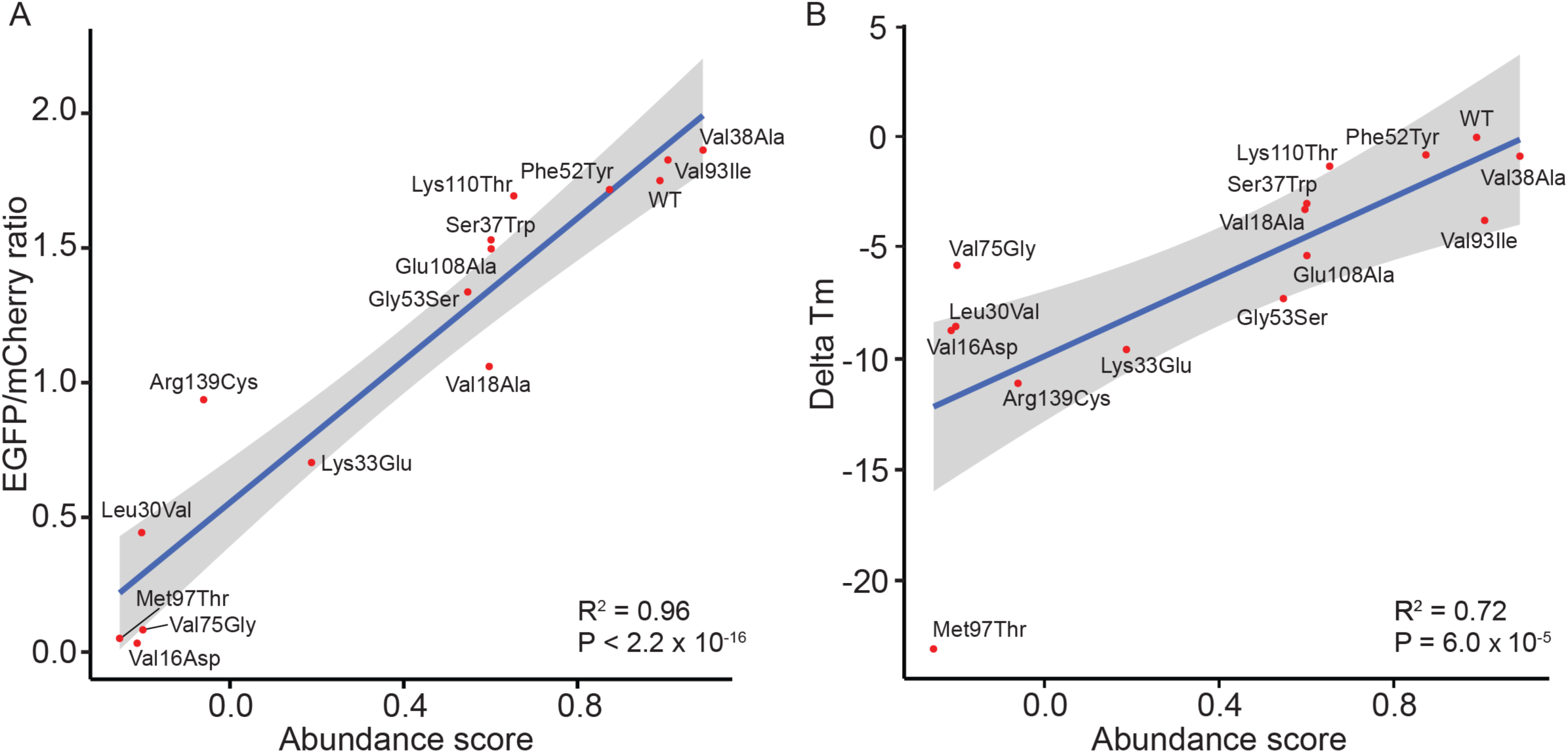
Orthogonal validation of *NUDT15* variant function. 14 *NUDT15* variants with a wide range of abundance scores were selected for validation. **A**) Fluorescence was measured in cells transfected with each *NUDT15* variant individually and plotted against estimated abundance score. **B**) Each of the 14 variant protein was expressed in *E coli*, purified to homogeneity, and subjected to thermostability assay. Melting temperature was determined using the Sypro®Orange and correlated with the abundance score calculated from the high throughput screen. P values were estimated using the Spearman test.

**Supplementary Fig 4.**
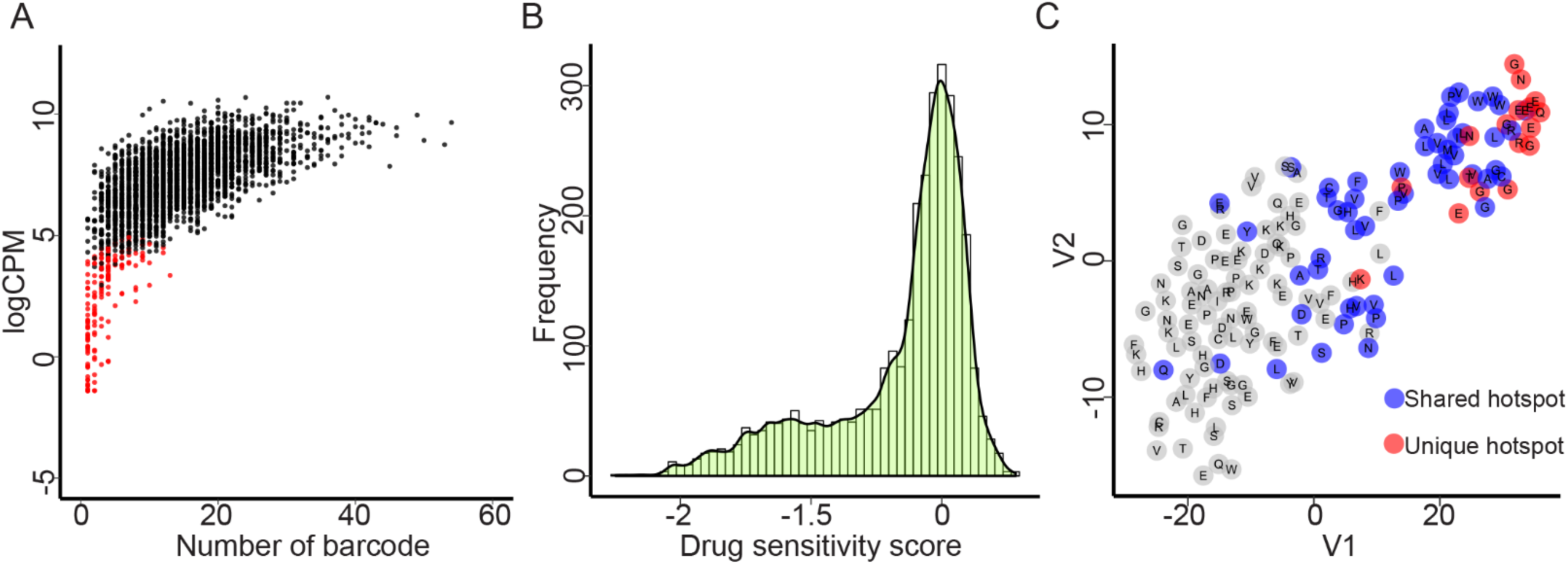
Drug sensitivity-based functional screen of *NUDT15* variants. **A**) Correlation between sequencing depth and the number of barcodes across 3,097 *NUDT15* variants, with each dot representing a unique variant. After quality control, 142 variants (red dots) were removed and 2,935 variants (black dots) remained for further analysis. **B**) Distribution of variant drug sensitivity score. **C**) t-SNE analysis identified residues at which genetic variation has similar effects on thiopurine cytotoxicity *in vitro*. Each dot represents a unique residue and clustering was on the basis of similarity and differences in the drug sensitivity score of 19 variants at each residue. Blue and red dots indicate residues as hotspots shared with abundance score-based assay or independently detected by drug screen-based assay, respectively. see **Methods**). Abbreviation. CPM, count per million

**Supplementary Fig 5.**
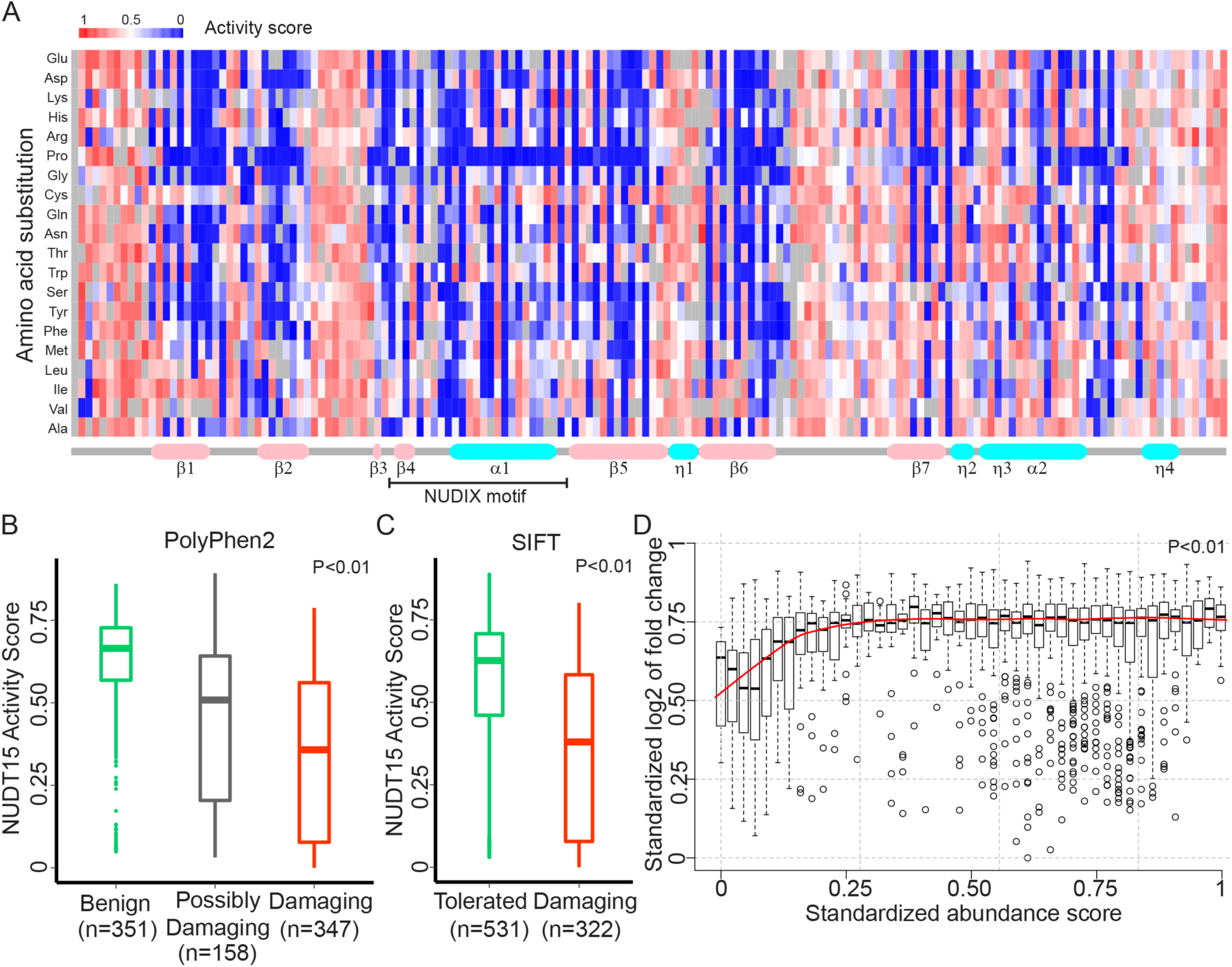
The development of NUDT15 activity score. **A**) Heatmap of NUDT15 activity score across 2,844 variants. Each column represents an amino acid residue in NUDT15 protein [from 1 to 164], and 19 rows indicate all 19 possible missense changes from the wildtype at this position. Red and blue denotes wildtype-like and damaging NUDT15 variants, respectively. Secondary structures are schematically indicated below the heatmap. Correlation between NUDT15 activity score with PolyPhen2 (**B**) and SIFT (**C**) predictions. Numbers represent variants classified in each category. P values were estimated by using the Rank Sum test. **D**) Correlation of abundance score and drug sensitivity score across *NUDT15* variant. P value was estimated by using the correlation test.

**Supplementary Fig 6.**
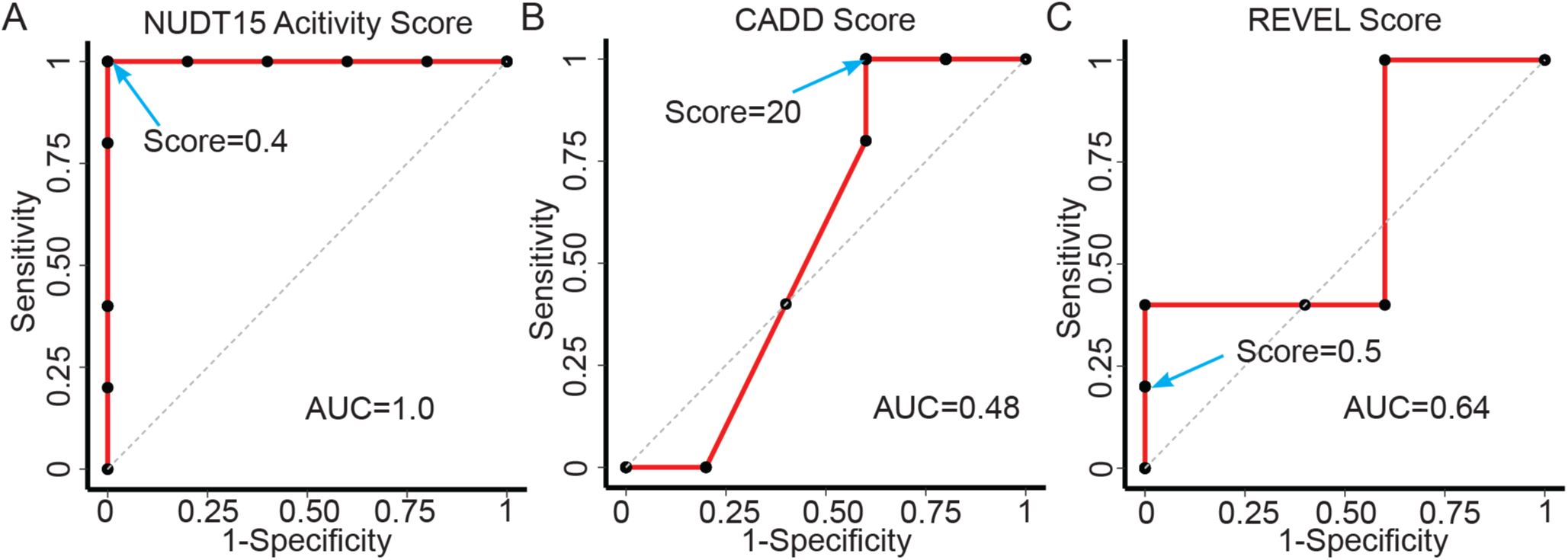
Receiver operating characteristic curves (ROC) analysis of *NUDT15* pharmacogenetic variants. ROC was plot for each score: A) NUDT15 activity score; B) CADD score; C) REVEL score. Proposed cutoff for each score is indicating by arrow. Analysis was based on 10 variants identified in 2,839 patients treated with thiopurine and NUDT15 pharmacogenetic variants were defined on the basis of clinical toxicity experienced by individuals who carried these variants (see **Supplementary Notes**).

**Supplementary Table 1.** Clinically observed *NUDT15* variants and their impact on thiopurine toxicity. 10 missense variants were identified by sequencing 2,398 patients, and were defined as either “toxicity-related” or “non-risk” (See **Methods**). For common variants, P values were estimated by using the linear regression model after adjusting for cohorts. Tolerated MP dosage in WT controls was either described in our previous publications *Nat Genet 48:367* and *J Clin Oncol 33:1235* (*), or estimated from ancestry-matched WT individuals treated on the same ALL protocol (^†^), with details included in **Supplementary Notes**. Abbreviations. NA, not applicable; sd, standard deviation; UK, United Kingdom; USA, United States of America; WT, wildtype.

| Variant | Cohort(s) | Case Number | Tolerated MP dosage (mg/m<sup>2</sup>),<br>mean (± sd) |  | Frequency | P value | Co-efficient | Final class |
| --- | --- | --- | --- | --- | --- | --- | --- | --- |
|  |  |  | Case | Ancestry-matched<br>WT control |  |  |  |  |
| Arg139Cys | USA | 27 | 50.5 (± 18.3) | 63.5 (± 14.8)* | Common | 4.0 × 10<sup>-9</sup> | -11.2 | Toxicity-related |
|  | Singapore | 21 | 32.7 (± 12.7) | 44.4 (± 13.3)* |  |  |  |  |
|  | Japan | 15 | 32.8 (± 21.8) | 45.9 (± 17.9)* |  |  |  |  |
|  | Guatemala | 13 | 29.3 (± 18.0) | 34.2 (± 14.4)* |  |  |  |  |
| Arg139His | USA | 5 | 52.6 (± 21.7) | 65.0 (± 13.6)* | Common | 0.029 | -9.5 | Toxicity-related |
|  | Guatemala | 6 | 27.1 (± 19.9) | 34.2 (± 14.4)* |  |  |  |  |
| Val18Ile | USA | 2 | 43.5 (± 5.5) | 60.0 (± 16.9)* | Common | 0.36 | -6.6 | Non-risk |
|  | Singapore | 2 | 42.0 (± 11.2) | 44.4 (± 13.3)* |  |  |  |  |
|  | Japan | 1 | 50.4 | 45.9 (± 17.9)* |  |  |  |  |
| Gln6Glu | USA | 1 | 58.3 | 60.0 (± 16.9)* | Rare | NA | NA | Non-risk |
| Arg11Gln | Hong Kong | 1 | 50.0 | 45.9 (± 14.9)† | Rare | NA | NA | Non-risk |
| Lys33Glu | UK | 1 | NA | NA | Rare | NA | NA | Toxicity-related |
| Arg34Thr | Singapore | 1 | 17.9 | 44.4 (± 13.3)* | Rare | NA | NA | Toxicity-related |
|  | Taiwan | 1 | 16.4 | 45.1 (± 15.3)† |  |  |  |  |
| Val75Glu | UK | 1 | NA | NA | Rare | NA | NA | Toxicity-related |
| Ser83Tyr | India | 1 | 52.9 | 49.7 (± 3.5)† | Rare | NA | NA | Non-risk |
| Val93Ile | India | 1 | 50.5 | 49.7 (± 3.5)† | Rare | NA | NA | Non-risk |

**Supplementary Table 3.**
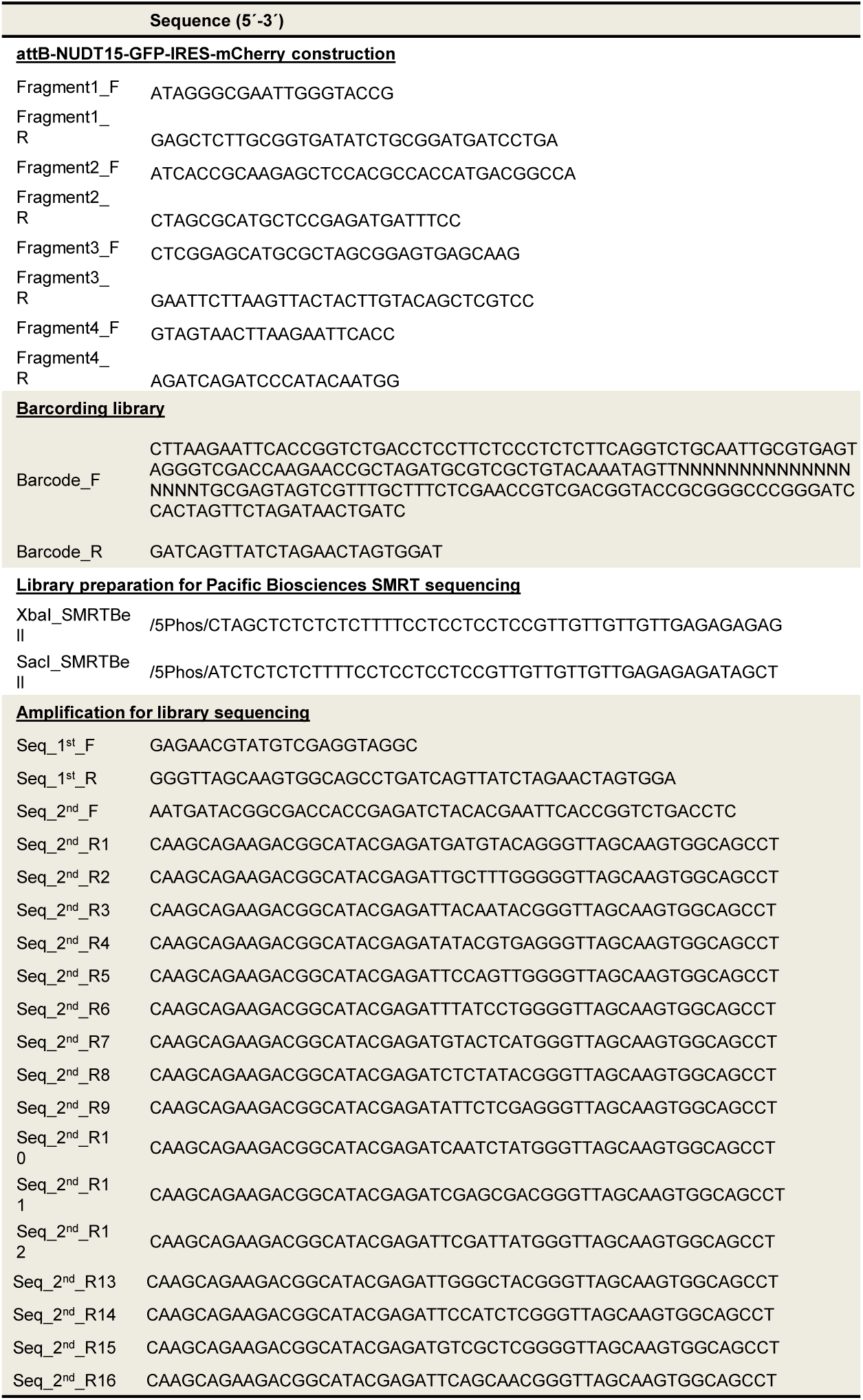
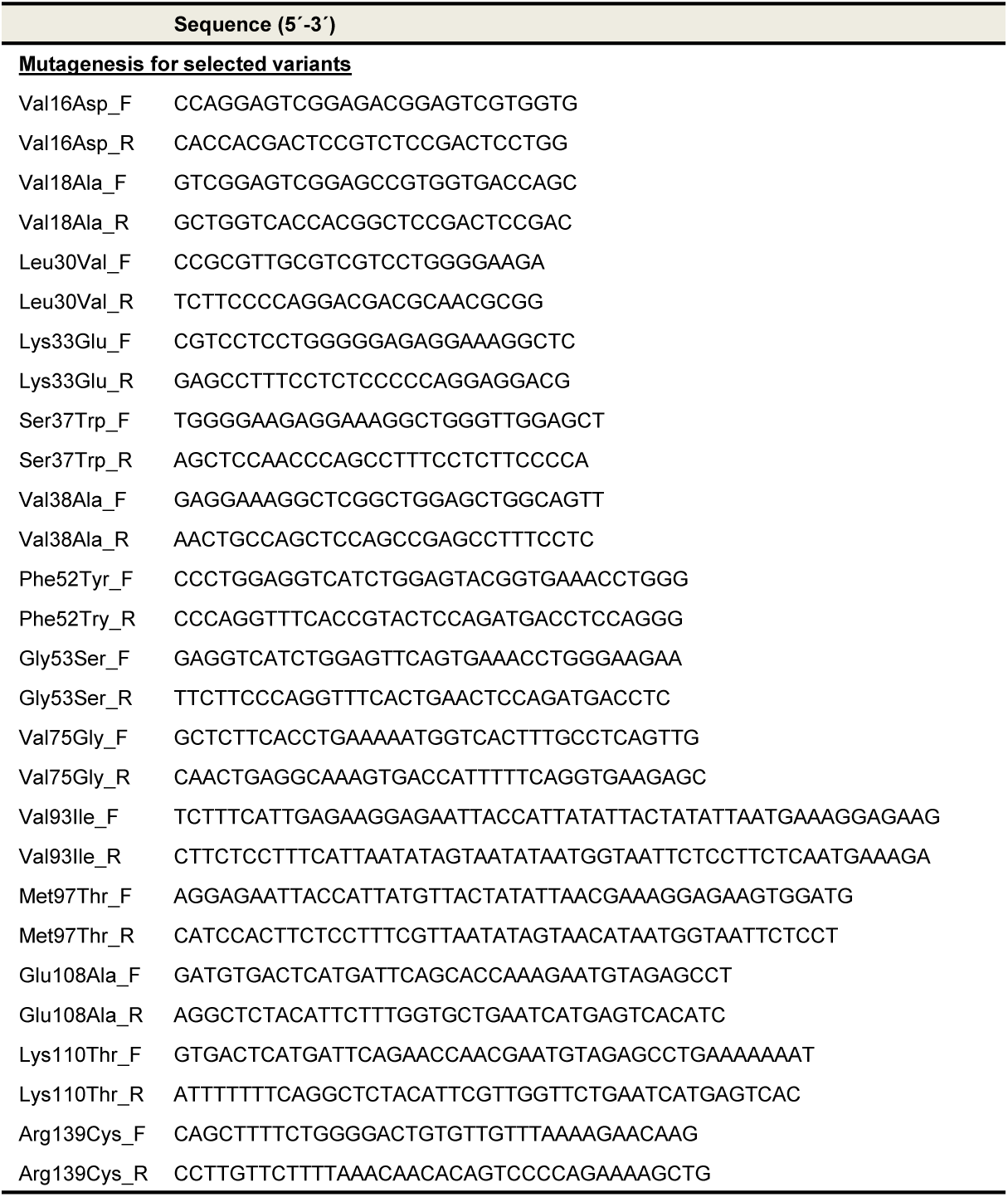
Primer sequences for library construction and cloning

**Supplementary Table 4.** Protocol-defined standard MP dosage and criteria for dose adjustment. ALL protocols in this study have a planned MP dosage ranging from 50 to 75 mg/m^2^ during maintenance therapy. MP dosage is clinically adjusted according to a target range of WBC and/or ANC. Abbreviations. ANC, absolute neutrophil count; HR, high risk; IR, intermediate risk; MP, 6-mercaptopurine; SR, standard risk; VHR, very-high risk; WBC, white blood cell; USA, United States of America.

| Cohort | Planned MP dosage (mg/m<sup>2</sup>) | Criteria for dose adjustment |
| --- | --- | --- |
| USA | 75 | ANC 0.75 – 1.5 x 10<sup>9</sup>/l |
| Singapore | 50 (SR, IR), 75 (HR) | WBC 2.0 – 4.0 x 10<sup>9</sup>/l |
| Japan | 50 | WBC 2.0 – 3.0 x 10<sup>9</sup>/l |
| Guatemala | 50 - 75 | WBC 1.5 – 3.0 x 10<sup>9</sup>/l |
| Taiwan | 60 (SR, IR), 75 (VHR) | WBC 1.8 – 3.0 x 10<sup>9</sup>/l<br>ANC 0.5 – 1.2 x 10<sup>9</sup>/l |
| Hong Kong | 50 | WBC 1.8 – 3.0 x 10<sup>9</sup>/l<br>ANC 0.5 – 1.2 x 10<sup>9</sup>/l |
| India | 60 | ANC 0.75 – 1.5 x 10<sup>9</sup>/l |

